# *In vivo* contribution of *Cyp24a1* promoter vitamin D response elements

**DOI:** 10.1101/2024.08.23.609393

**Authors:** Mark B. Meyer, Seong Min Lee, Jordan M. Towne, Shannon R. Cichanski, Martin Kaufmann, Glenville Jones, J. Wesley Pike

## Abstract

CYP24A1 is a multifunctional, P450 mitochondrial 24-hydroxylase enzyme that is responsible for catabolism of the most active vitamin D hormone (calcitriol, 1,25(OH)_2_D_3_), its precursor (calcifediol, 25(OH)D_3_), and numerous other vitamin D metabolites at the 23- and 24-carbon positions. In the kidney, *Cyp24a1* is induced by 1,25(OH)_2_D_3_, induced by FGF23, and potently suppressed by PTH to tightly control the circulating blood levels of 1,25(OH)_2_D_3_. This gene is believed to be under the control of a pair of classic promoter proximal (PRO) vitamin D response elements (VDREs) that are aided by distal, downstream (DS) containing enhancers that we identified more recently. The DS1 enhancer cluster was found to respond to PTH and FGF23 actions in a kidney-specific manner. The DS2 enhancer cluster was found to assist in the response of 1,25(OH)_2_D_3_ in kidney, as well as other target tissues. Despite this knowledge, the *in vivo* contribution of the PRO VDREs to gene expression, what drives *Cyp24a1* basal expression in the kidney, how FGF23 activates *Cyp24a1*, and importantly, how PTH suppresses *Cyp24a1*, all remain unknown. Here in this study, we utilize homology directed CRISPR to mutate one or both VDREs in the PRO region of the *Cyp24a1* gene *in vivo* in the mouse to address these questions. We found that the VDRE (VDRE1) more proximal to the to the transcriptional start site (TSS) is the dominant VDRE of the pair and mutation of both VDREs leads to a dramatic loss of VDR, a reduction of *Cyp24a1* gene expression in the kidney, and a near elimination of 1,25(OH)_2_D_3_ induction in the intestine. FGF23 induction of *Cyp24a1* was reduced with mutation of the PRO VDREs, however, co-treatment of 1,25(OH)_2_D_3_ and FGF23 synergistically increased *Cyp24a1* expression even with the loss of the PRO VDREs. PTH suppression of *Cyp24a1* gene expression was unchanged with PRO VDRE mutations, despite a minor reduction in total pCREB occupancy. Finally, VDR occupancy was dramatically reduced across the DS enhancers in the *Cyp24a1* locus after the PRO VDREs mutation. Taken together, our data suggest a cooperative relationship between the DS and PRO enhancers in the regulation of *Cyp24a1* by 1,25(OH)_2_D_3_ and FGF23, and despite the overall reduction of CREB on the genome it appeared that suppression either does not rely on CREB or that the PRO VDREs are unconnected to PTH suppression altogether. These studies point to the DS1 region as a basal switch for *Cyp24a1* expression and help further define the interconnected genomic control of these hormones on vitamin D catabolism.

## Introduction

Calcium (Ca) and phosphate (P) homeostasis is tightly controlled by regulation of the most active vitamin D metabolite, 1,25(OH)_2_D_3_ (1α,25-dihydroxyvitamin D_3_, calcitriol) (1). Dietary or UV-dependent vitamin D_3_ is converted in the liver to 25(OH)D_3_, which is then further hydroxylated at the 1-alpha position by the CYP27B1 (1α-hydroxylase) enzyme to create 1,25(OH)_2_D_3_ (2-4). 1,25(OH)_2_D_3_ is then inactivated and marked for degradation by the 24-hydroxylase enzyme, CYP24A1 (5). CYP24A1 is a multifunctional enzyme that continues to hydroxylate and modify vitamin D metabolites at both the carbon 24 and 23 positions further down the degradation pathway (6,7). The expression of these enzymes is reciprocally controlled by parathyroid hormone (PTH), fibroblast growth factor 23 (FGF23), and 1,25(OH)_2_D_3_ itself, in the kidney (8). PTH induces expression of the *Cyp27b1* gene, and potently suppresses the *Cyp24a1* gene, whereas 1,25(OH)_2_D_3_ and FGF23 both suppress *Cyp27b1* and induce *Cyp24a1* (8). Outside of the kidney, both genes have very little basal expression and are not controlled by these endocrine hormones due to kidney-specific enhancers for both *Cyp27b1* and *Cyp24a1* (9,10).

The *Cyp24a1* gene is highly responsive to 1,25(OH)_2_D_3_ concentrations in the blood and is known to be regulated by the 1,25(OH)_2_D_3_-bound vitamin D receptor (VDR). A pair of vitamin D response elements (VDREs) were found upstream of *Cyp24a1* upstream from the transcriptional start site (TSS) at ∼ -150 and -250 bp (11-13). The sequences surrounding these VDREs were further examined using reporter constructs to determine their activity and contributions (12,14-17). Using large-scale genome-wide ChIP, we discovered that the regulation of *Cyp24a1* was aided by downstream enhancers as well (18), and in fact, much of the genomic binding of VDR was found to be in inter- and intra-genic regions (19,20). In further studies, we defined these downstream (DS) enhancers at *Cyp24a1* and their activities in both kidney and non-renal tissues *in vivo* (10). In addition to the promoter (PRO) region (+10 to -350 bp) of *Cyp24a1*, we found the downstream 1 (DS1, +21 kb to +32 kb) region controlled *Cyp24a1* expression via PTH and FGF23 in kidney, and the downstream 2 (DS2, +35 kb to +46 kb) region controlled *Cyp24a1* expression via 1,25(OH)_2_D_3_ in all tissues. The DS1 set of enhancers were only present in the kidney and not in other tissues (8). Importantly, the basal level of *Cyp24a1* is highest in the kidney and virtually not expressed in many tissues (8,10). Only the DS1 deletion reduced the baseline activity of *Cyp24a1* in the kidney and DS1 deletion also greatly reduced the effects of PTH or FGF23 on *Cyp24a1* expression (10). Since 1,25(OH)_2_D_3_ induction of *Cyp24a1* is rather ubiquitous across tissues, we concluded that serum PTH and FGF23 are likely more important than 1,25(OH)_2_D_3_ for maintaining basal expression in the kidney (10).

The mechanisms of *Cyp24a1* regulation by PTH and FGF23 continue to remain largely unknown. From our recent work, we found that while CREB was highly recruited for activation of *Cyp27b1*, CREB was unchanged in occupancy during PTH suppression of *Cyp24a1* (21). However, many associated coactivators like CRTC2, CBP, SRC-1, and others, were greatly reduced across the locus (21). CREB did appear bound to the *Cyp24a1* promoter and although CREB response elements (CREs) can be found *in silico*, the relationship of CREB to *Cyp24a1* activity at the promoter has not been tested. We hypothesized that CREB may be recruiting histone deacetylase complexes to complete this suppression, however, this mechanism is not well-defined (21). FGF23 activation of *Cyp24a1* has remained a complete enigma due to the gap of knowledge between FGF23 signaling and the identity of the signal-related transcription factor(s) (TF) involved for FGF23-mediated genomic activation. Despite the lack of FGF23-responsive TF, we know that FGF23 is an activator of *Cyp24a1* like 1,25(OH)_2_D_3_ and shares many of the same histone markers and coactivators recruited during activation (8,21).

Here in this study, we examined the relationship between VDR binding to the VDREs in the PRO region and the regulation of *Cyp24a1* activation, suppression, and baseline. We hypothesized that reductions of VDR near the promoter would destabilize the downstream enhancer structure of both DS1 and DS2 leading to a reduction of *Cyp24a1* activity and a dysregulation of vitamin D metabolism. Through these mutations, we found that VDR binding was indeed greatly compromised which led to reductions of *Cyp24a1* activity by 1,25(OH)_2_D_3_. However, we also found an impairment of FGF23-mediated activity of *Cyp24a1*. While PTH suppression was unchanged, we also found a decrease of CREB binding across the locus. Finally, the mineral phenotype of the mutated mice were unremarkable from WT and there were minor changes to vitamin D metabolites in the blood. Taken together, our data demonstrate that the PRO VDREs was required for full activation, however, mineral homeostasis and basal activity was not dependent on the activities of *Cyp24a1* induction from the PRO VDREs in the mouse.

## Results

### Reduction in 1,25(OH)_2_D_3_-mediated induction of Cyp24a1

Our prior studies hypothesized that the DS regions (DS1 and DS2) worked together with the PRO region and perhaps specifically, the PRO VDREs to control the expression of *Cyp24a1 in vivo* (18,22). This association was implied by the activities that were reduced, yet retained, during the deletion of the DS1 and DS2 regions (22). At that time, this relationship was not explicitly tested given the complexities of the promoter region sequence. For this current study, we devised a mutation strategy that specifically mutated the VDRE sequences in the mouse at both the -156 to -171 bp (VDRE1) and -267 to -282 bp (VDRE2) upstream from the transcriptional start site (TSS) as detailed in Fig. 1A. There has been considerable debate around an AP-1 sequence that may contribute to the activation *in vivo* as it is adjacent to VDRE1 (12,16,23). This AP-1 sequence, GAGTCA, is similar sequence to the consensus VDRE half site (G/A GGT G/C A) and is positioned 3 nucleotides, the appropriate spacing, from the accepted VDRE1 sequence (AGGTCAgtgAGGGCG) (24-26). To remove the possibility of activation of this sequence, we mutated the AP-1 site as well (Fig. 1B). In addition to the AP-1 site, an ETS binding site (EBS) was also annotated adjacent to the AP-1 – VDRE1 sequence (16). The core of this ETS binding site was not mutated in our mouse. Homology directed recombination (HDR) CRISPR/Cas9 was utilized to make this mutation *in vivo* with a long, single stranded donor oligo (ssODN) injected with dual guide RNAs targeting the PRO region in C57BL/6J mice. From this injection, we obtained several germline founder mice with the exact desired mutation, as well as a mouse with mutation only in the VDRE2 sequence. We named the double mutation (VDRE1/AP-1 and VDRE2) mouse the “C24-V1V2” mouse and the single VDRE2 mutation (WT sequence for VDRE1/AP-1) mouse the “C24-V2” mouse.

**Figure 1:**
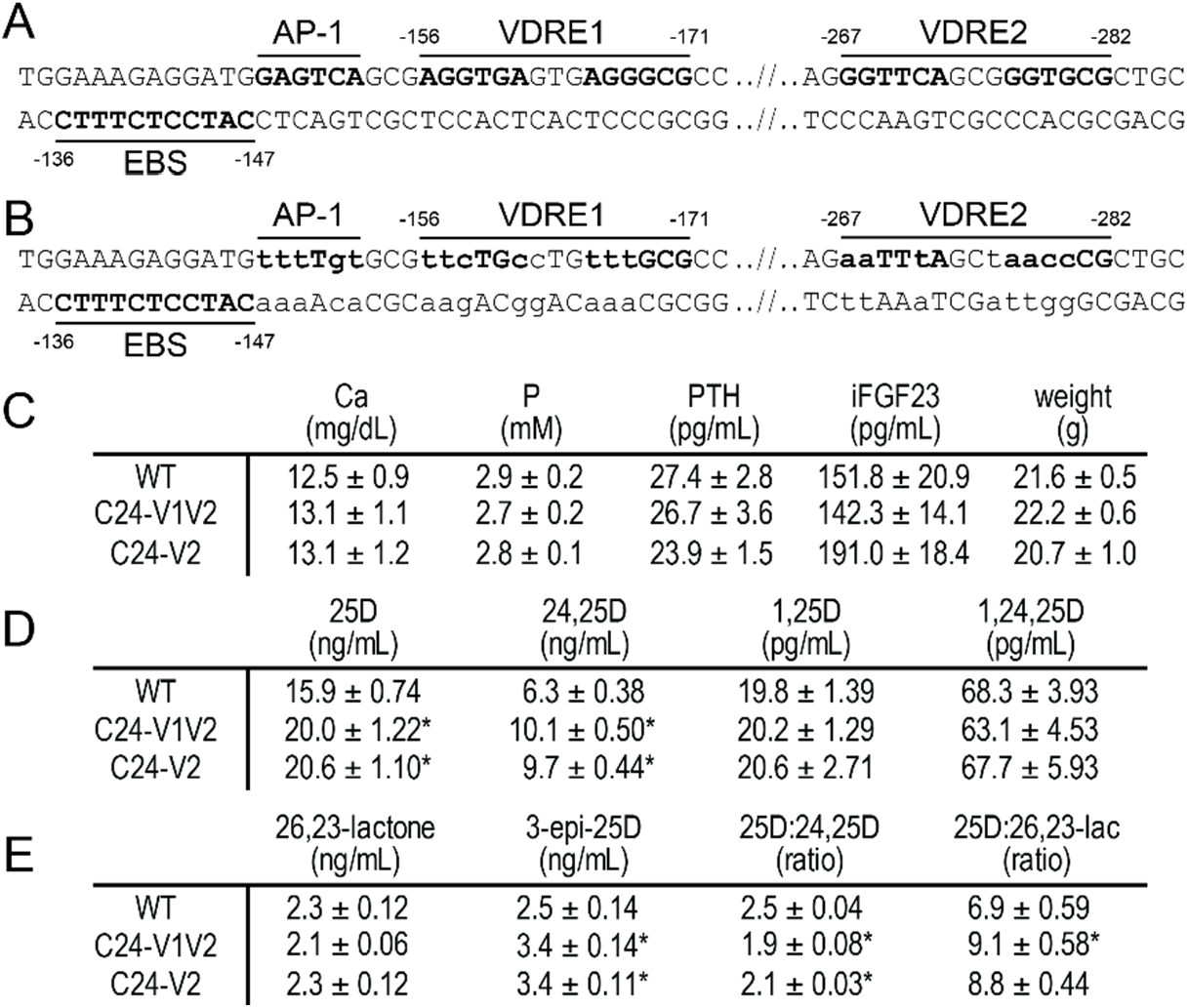
C24-V1V2 and C24-V2 mutations and basal phenotyping. Sequences both WT (A) and mutated (B) in the *Cyp24a1* promoter region with distances indicated from the transcriptional start site. Elements are labeled above or below DNA sequence and mutated sequences indicated in lower case. C, serum measurements for calcium (Ca), phosphate (P), PTH, in-tact FGF23 (iFGF23), and body weight of the C24-V1V2, C24-V2, and their wildtype (WT) littermates. Vitamin D metabolite profile for 25(OH)D_3_, 24,25(OH)_2_D_3_, 1,25(OH)_2_D_3_, and 1,24,25(OH)_3_D_3_ (A) and 25(OH)D_3_-26,23-lactone (26,23-lactone), 3-epi-25(OH)D_3_ (3-epi-25D) and the ratios of 25(OH)D_3_:24,25(OH)_2_D_3_ and 25(OH)D_3_:25(OH)D_3_-26,23-lactone (B). All values displayed as averaged values ± SEM. WT, n=10; C24-V1V2, n=6; C24-V2, n=6. Statistics performed by one-way ANOVA with ^*^, p < 0.05, genotype vs WT.

Basal phenotyping was performed in both the C24-V1V2, C24-V2 mice, and their WT littermates, including an analysis of the vitamin D metabolites (8-10 weeks of age). The serum calcium (Ca), phosphate (P), in-tact FGF23 (iFGF23), PTH, and weights were all not significantly different than their WT littermates (Fig. 1C) and were in line with prior measurements in our WT animals (9,22). Despite no apparent changes in the Ca/P homeostatic system, we hypothesize that the potential for *Cyp24a1* activity reduction after VDREs deletion could result in altered vitamin D metabolic profile. Examining a wide panel of vitamin D metabolites, we measured 25(OH)D_3_, 24,25(OH)D_3_, 1,25(OH)_2_D_3_, and 1,24,25(OH)_2_D_3_ in Fig. 1D. There was a slight elevation in the 25(OH)D_3_ levels of both the C24-V1V2 and C24-V2 mice compared to their WT counterparts Fig. 1D, however this was very mild compared to the *Cyp24a1* and *Cyp27b1* knockout animals (9,22). Additionally, the mice had elevated levels of 24,25(OH)_2_D_3_, the catabolic product of 25(OH)D_3_ that is processed by the CYP24A1 enzyme. Other catabolic products from CYP24A1 are the 25(OH)D_3_-26,23-lactone and 3-epi-25(OH)D_3_ metabolites and these, along with the ratios of 25(OH)D_3_:24,25(OH)_2_D_3_ and 25(OH)D_3_:26,23-lactone are shown in Fig. 1E. These metabolites are more reliable markers of CYP24A1 activity because they are in less flux than 24,25(OH)_2_D_3_ measurement alone (27-29). In these C24-V1V2 and C24-V2 mice, the 25(OH)D_3_-26,23-lactone was unchanged from WT, however the 3-epi-25(OH)D_3_ was elevated (Fig. 1E). This also resulted in a decreased ratio of 25(OH)D_3_:24,25(OH)D_3_ and 25(OH)D_3_:26,23-lactone as can be seen in Fig. 1E. The accumulation of 25(OH)D_3_, the substrate for both CYP24A1 and CYP27B1, may indicate an overall decrease of CYP24A1 or CYP27B1 enzymatic activity or loss of expression, so we next examined the gene expression in these mouse models.

The C24-V1V2, C24-V2, and WT littermates were injected with 10 ng/g bw 1,25(OH)_2_D_3_ for 6 h and gene expression for *Cyp24a1, Cyp27b1*, and *Vdr* was examined in both the kidney and intestine tissue (Fig. 2). The 1,25(OH)_2_D_3_ induction of *Cyp24a1* was reduced ∼ 60% in the kidney (Fig. 2A). *Cyp27b1* suppression by 1,25(OH)_2_D_3_ in the kidney appeared unchanged (Fig. 2B), however, the baseline levels of *Cyp27b1* were reduced ∼ 50%. The *Vdr* levels in the kidney (Fig. 2C) of the C24-V1V2 mice, both basal and induced, were unchanged. The C24-V2 mice, however, only had a 10% reduction in 1,25(OH)_2_D_3_-induced activity of *Cyp24a1* in the kidney (Fig. 2A), they retained their 1,25(OH)_2_D_3_-mediated suppression of *Cyp27b1* (Fig. 2B), and the *Vdr* levels were unchanged as well (Fig. 2C). Despite the normal serum Ca, P, PTH, and iFGF23, minor fluctuations may be causing an expression imbalance that is leading to the mildly decreased *Cyp27b1* expression, but seemingly normal basal *Cyp24a1* expression.

**Figure 2:**
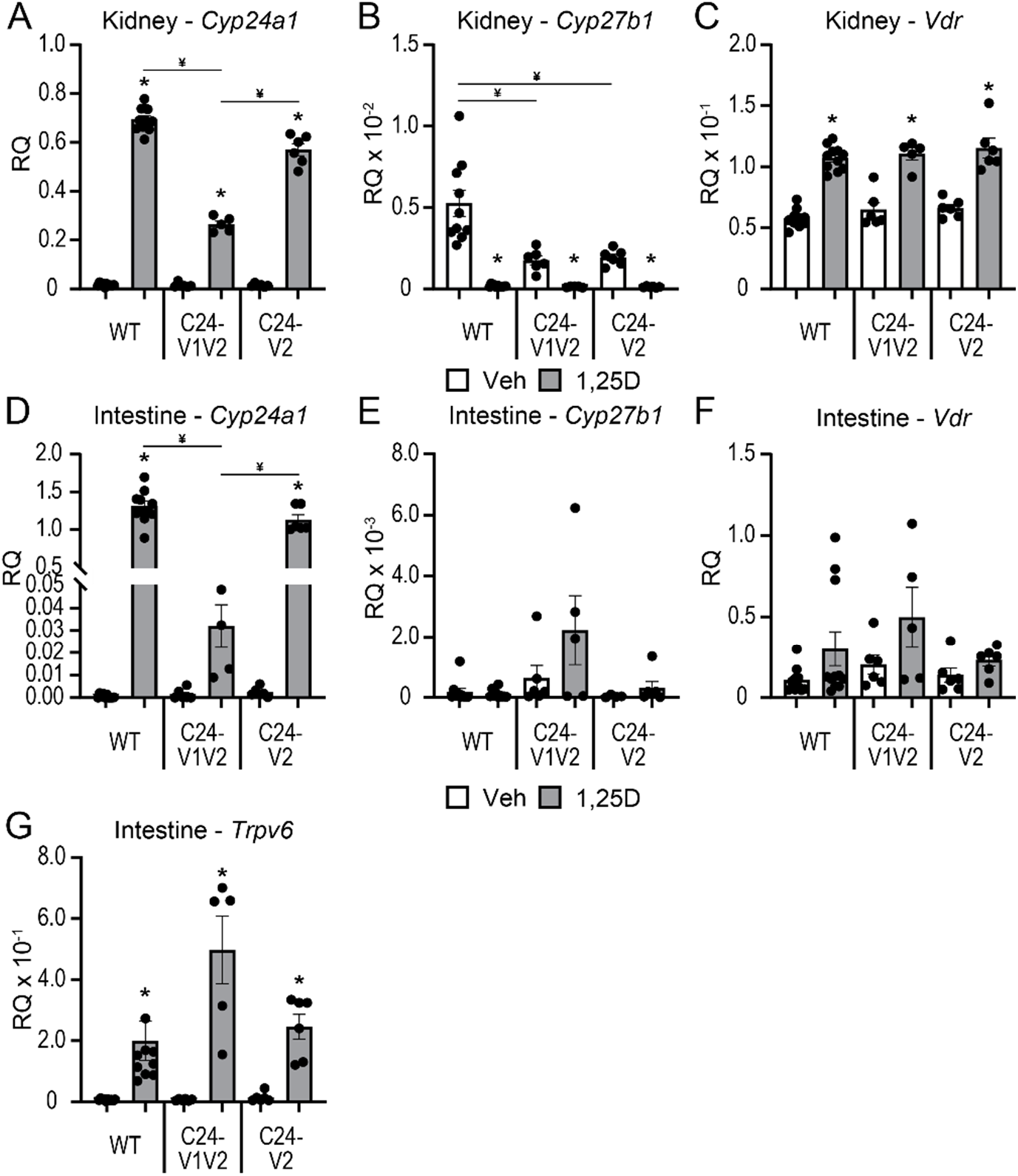
1,25(OH)_2_D_3_-mediated gene expression changes in mutant mice. Gene expression was performed in WT, C24-V1V2, and C24-V2 mice injected with 10 ng/g bw 1,25(OH)_2_D_3_ (1,25D) or ethanol vehicle (Veh) for 6 h for kidney *Cyp24a1* (A), *Cyp27b1* (B), and *Vdr* (C) and intestine *Cyp24a1* (D), *Cyp27b1* (E), *Vdr* (F), and *Trpv6* (G). Data are displayed as relative quantitation (RQ) against *Gapdh*. WT (Veh, 1,25D) n ≥ 10; C24-V1V2 (Veh, 1,25D), n ≥ 6; C24-V2 (Veh, 1,25D), n ≥ 6. 2-way ANOVA with tukey post-tests comparing both treatment and genotype was performed. ^*^, p < 0.05, 1,25(OH)_2_D_3_ vs Veh; ¥, p < 0.05, selected pairs as indicated.

The chromatin architecture, gene expression, and DS enhancers are different between the kidney and nonrenal tissues as we have extensively reported (8,10,30), and the DS2 region, but not the DS1 region, had an impact on these tissues such as the intestine (10,18,31). To test the impact of the PRO VDREs on nonrenal tissues, intestinal (duodenal) *Cyp24a1, Cyp27b1*, and *Vdr* expression were measured in C24-V1V2, C24-V2, and WT littermate mice injected with 1,25(OH)_2_D_3_. Here, the 1,25(OH)_2_D_3_-mediated *Cyp24a1* induction was reduced by > 90% in the C24-V1V2 mice, and the C24-V2 mice were not statistically different than WT 1,25(OH)_2_D_3_-mediated induction (Fig. 2D). *Cyp27b1* has very low expression in the intestine, and it is not regulated by 1,25(OH)_2_D_3_, so therefore was unaffected (Fig. 2E). *Vdr* expression was unchanged as well in either mouse model (Fig. 2F). We also examined the duodenal expression of the calcium channel and vitamin D responsive gene, *Trpv6*, to ensure that calcium transport was not altered in the C24-V1V2 and C24-V2 mice. The baseline and induction of *Trpv6* were unchanged (Fig. 3G). Finally, we conducted a 1,25(OH)_2_D_3_ time course (0, 3, 6, 12, 24 h) in both the kidney and intestine of the C24-V1V2 and C24-V2 mice (Fig. 3). These data confirm our single time point data and do not display any altered kinetics of 1,25(OH)_2_D_3_ response in the mutant mice. While these mice have dramatic changes in 1,25(OH)_2_D_3_-mediated challenges, it appears they retain a normal phenotype and mineral homeostasis through alterations of vitamin D metabolites without the VDREs in the PRO region of *Cyp24a1*.

**Figure 3:**
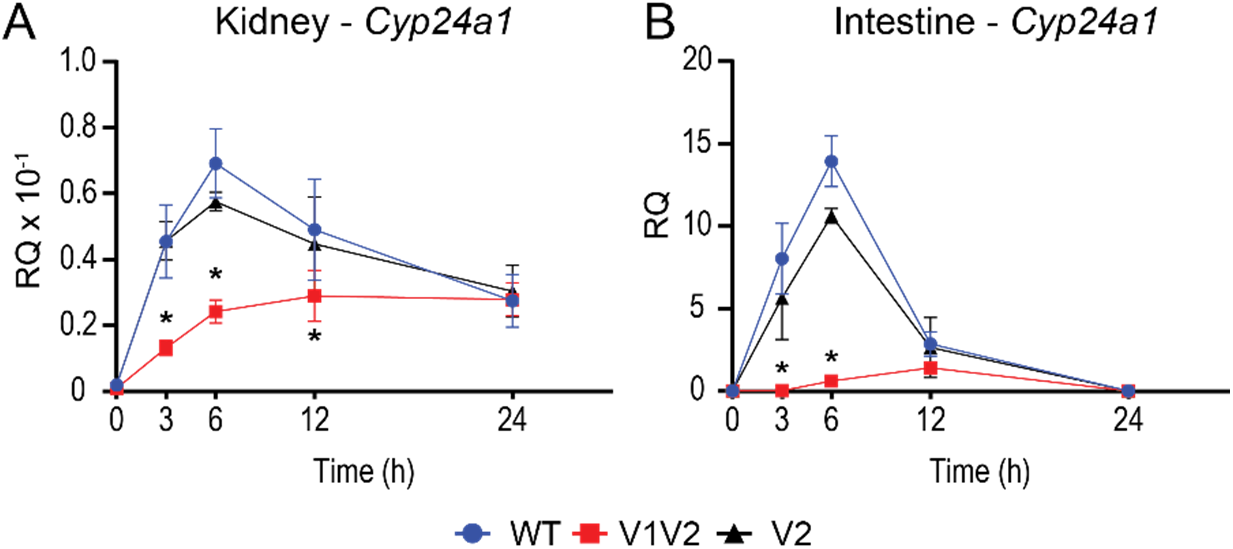
Time course activation by 1,25(OH)_2_D_3_ in mice lacking *Cyp24a1* PRO VDREs. Gene expression was performed in WT, C24-V1V2, and C24-V2 mice injected with 10 ng/g bw 1,25(OH)_2_D_3_ (1,25D) or ethanol vehicle (Veh) for 0, 3, 6, 12, or 24 h for kidney *Cyp24a1* (A) and intestine *Cyp24a1* (B). Data are displayed as relative quantitation (RQ) against *Gapdh*. WT, C24V1V2, C24V2, n = 6. 2-way ANOVA with tukey post-tests comparing both time and genotype was performed. ^*^, p < 0.05, genotype vs WT within each time point.

### Associated activities of FGF23 and PTH in the VDRE mutated mice

In addition to 1,25(OH)_2_D_3_, FGF23 and PTH are potent modulators of *Cyp24a1* expression and activity in the kidney. As mentioned, these actions are unique due to the presence of tissue specific enhancers located in the DS2 region (10,30). Despite our characterization of the DS1 region, it is unknown how the promoter region coordinates with the DS1 enhancer and furthermore we do not understand if this complex involves VDR or is independent of VDR. To investigate these possibilities, we injected our C24-V1V2, C24-V2, or WT littermate mice with 50 ng/g bw recombinant FGF23 (rFGF23) for 3 h, 230 ng/g bw PTH for 1 h, and compared to PBS vehicle. We then examined the expression of *Cyp24a1, Cyp27b1*, and *Vdr* in the kidney as well as the intestine. As can be seen in Fig. 4A, FGF23 induced while PTH potently suppressed *Cyp24a1* expression in the kidney in the WT mice. As observed in the 1,25(OH)_2_D_3_ experiments (Fig. 2), the basal expression of *Cyp24a1* was unchanged in the C24-V1V2 or C24-V2 mice. Additionally, PTH suppression of *Cyp24a1* was also unchanged in these mice compared to their WT littermates. FGF23 induction of *Cyp24a1*, on the other hand, was significantly reduced (∼ 50%) in the C24-V1V2 and the C24-V2 mice (Fig. 4A). *Cyp27b1* induction by PTH appeared to be reduced in the C24-V1V2 model (Fig. 4B), however the baseline expression of *Cyp27b1* was reduced as observed in Fig. 1 and thus, the fold change of induction was unchanged. FGF23 suppression of *Cyp27b1* was unaltered between genotypes. Finally, *Vdr* expression was unchanged in the kidney as well in vehicle treated mice (Fig. 4C). There was a reduction of *Vdr* with FGF23 treatment in the C24-V2 mice, however this appears to be inconsistent with the other models. We also examined these genes in the intestine, however, FGF23 and PTH do not regulate *Cyp27b1* or *Cyp24a1* in the intestine and therefore no change was observed for these hormones on the selected genes (Fig. 4D-F). Overall, based on these results, PTH suppression of *Cyp24a1* in the kidney does not appear to require the PRO VDREs, however, FGF23 induction was reduced in both models. This implicates a relationship between VDR/1,25(OH)D_3_ and FGF23 and/or the unknown transcription factor through which FGF23 activates the genome.

**Figure 4:**
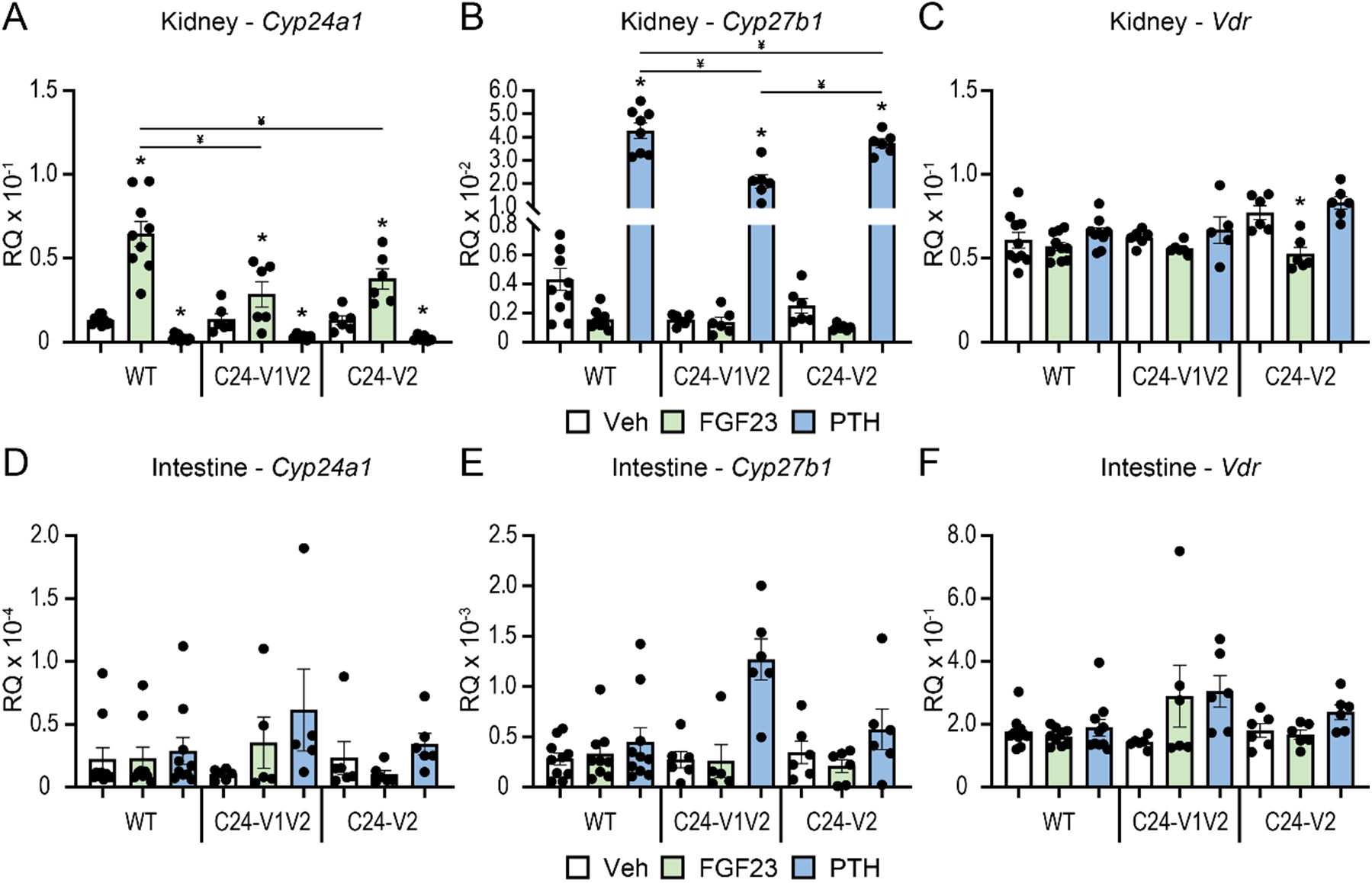
Contribution of PRO VDREs to FGF23- and PTH-mediated *Cyp24a1* regulation. Gene expression was performed in WT, C24-V1V2, and C24-V2 mice injected with 230 ng/g bw PTH (1 h), 50 ng/g bw FGF23 (3 h) or PBS vehicle (Veh) for kidney *Cyp24a1* (A), *Cyp27b1* (B), and *Vdr* (C) and intestine *Cyp24a1* (D), *Cyp27b1* (E), and *Vdr* (F). Data are displayed as relative quantitation (RQ) against *Gapdh*. WT (Veh, FGF23, PTH) n ≥ 10; C24-V1V2 (Veh, FGF23, PTH), n ≥ 6; C24-V2 (Veh, FGF23, PTH), n ≥ 6. 2-way ANOVA with tukey post-tests comparing both treatment and genotype was performed. ^*^, p < 0.05, treatment vs Veh; ¥, p < 0.05, selected pairs as indicated.

### Genomic changes of transcription factor occupancy without the PRO VDREs

Based on the gene expression changes and to confirm reduction of VDR binding in the promoter with mutated VDREs, we next examined genomic binding of VDR after 1,25(OH)_2_D_3_ treatment by ChIP-seq across the *Cyp24a1* locus. The PRO, DS1, and DS2 regions were previously all found to bind VDR in the kidney, whereas the PRO and DS2 region were only occupied by VDR in the intestine (10). To examine these mutated regions, a new reference genome was built for mm9 (mm9-C24m) that contained the mutations as described in Fig. 1B for the *Cyp24a1* PRO region. The mutations were otherwise seamless as confirmed by deep sequencing, therefore the genomic coordinates and positioning were unchanged for the mm9 reference outside of these regions. ChIP was performed in both the kidney and intestine (duodenum) of WT and C24-V1V2 mice after treatment with 1,25(OH)_2_D_3_ for 1 h (9,10,22). Given the rather mild phenotype in the C24-V2 mice by gene expression, ChIP-seq was not performed in these animals. Data from the WT littermates (upper tracks) match our previous data for the *Cyp24a1* locus as recently published for VDR in both the kidney (Fig. 5A) and the intestine (Fig. 5B) (10,21). The lower tracks are data from the C24-V1V2 mice. For the kidney (Fig. 5A), we found that the VDR binding was reduced (reduced peak height) across the entire genomic locus and furthermore, there was a reduction in the increased occupancy by VDR after 1,25(OH)_2_D_3_ treatment (decrease of peak height, blue color). At the PRO region, a very small amount of VDR was found recruited in the absence of the VDREs, however, there was no 1,25(OH)_2_D_3_-induced increased occupancy in this region. In the intestine (Fig. 5B), we found that C24-V1V2 had further decreases of VDR binding compared with the kidney occupancy. As expected, VDR binding in the DS1 region is absent from the intestine. These activities were specific to *Cyp24a1* as ChIP-seq data in C24-V1V2 mice for *Cyp27b1, Vdr*, and *Trpv6* were all unchanged from WT (Fig. 6) at previously annotated enhancers (22,32,33).

**Figure 5:**
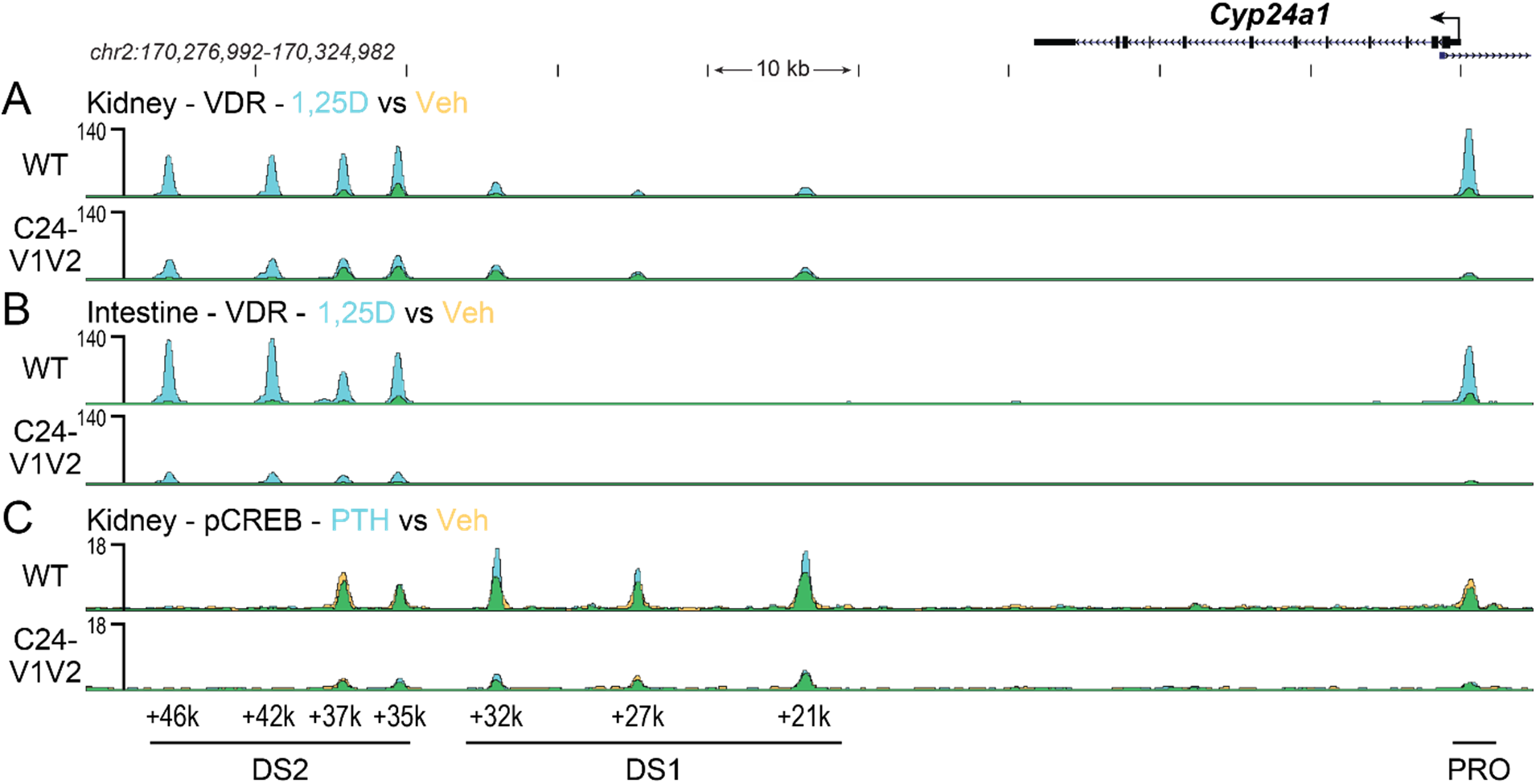
Genomic occupancy of VDR and CREB in the *Cyp24a1* locus. ChIP-seq analysis near *Cyp24a1* for kidney VDR (A) and intestine VDR (B) from WT or C24-V1V2 mice injected with 10 ng/g bw 1,25(OH)_2_D_3_ for 1 h (n=3). C, ChIP-seq for kidney phosphorylated CREB (pCREB) from WT or C24-V1V2 mice injected with 230 ng/g bw PTH for 30 mins (n=3). Overlaid triplicate and averaged ChIP-seq tracks where vehicle (Veh) are shown in yellow, treatments shown in blue, and overlapping data appear as green. Regions of interest are denoted at the bottom of the figure (PRO, DS1, or DS2). Genomic region displayed is chr2:170,276,992-170,324,982 and maximum height of tag sequence density for each data track indicated on the Y-axis (normalized to input and 10^7^ tags).

**Figure 6:**
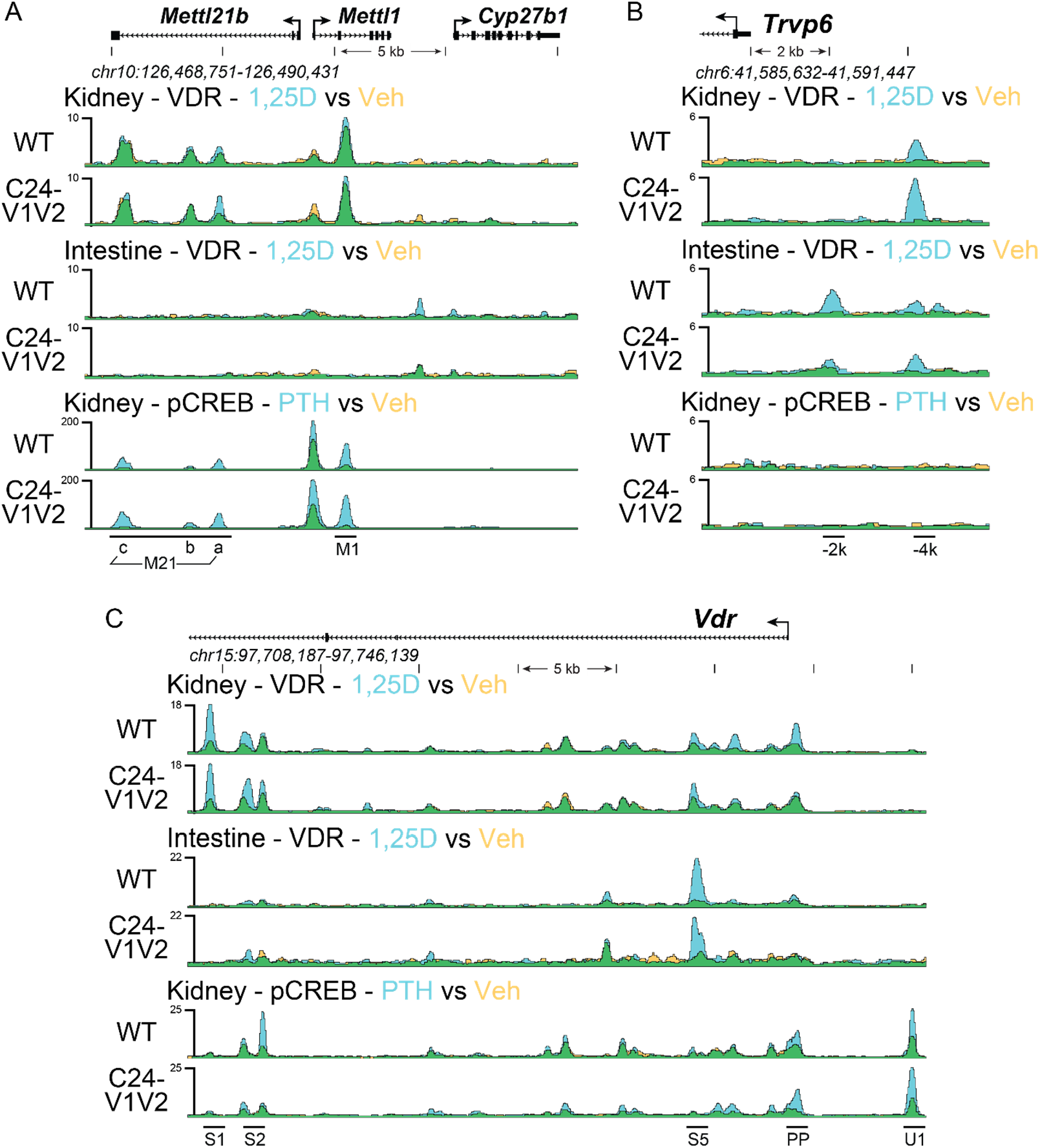
Genomic occupancy near *Cyp27b1, Vdr*, and *Trpv6*. ChIP-seq analysis near *Cyp27b1, Vdr*, and *Trpv6* for kidney and intestine as in Fig. 5. Overlaid triplicate and averaged ChIP-seq tracks where vehicle (Veh) are shown in yellow, treatments shown in blue, and overlapping data appear as green. Regions of interest are denoted at the bottom of the figure from prior studies (22,32,33). Genomic regions displayed are indicated for each region and maximum height of tag sequence density for each data track indicated on the Y-axis (normalized to input and 10^7^ tags).

PTH is a potent suppressor of *Cyp24a1* in the kidney and not in other tissues in the body (9,10,21,22), however, we do not yet understand the mechanism through which this occurs. Our recent work demonstrated that pCREB was largely unchanged despite reductions in coactivators (21). We also observed that pCREB was recruited to the PRO region of *Cyp24a1*, but unaltered by PTH treatment (21). In Fig. 4, we found that PTH suppression of *Cyp24a1* gene expression in C24-V1V2 mice was unchanged. To test a connection between the PRO VDREs and DS1 in the activity of PTH suppression, we performed ChIP-seq with pCREB after 230 ng/g bw injection of PTH or vehicle (Veh) for 30 mins. As seen in Fig. 5C (upper track), the WT mice recruited pCREB to the PRO, DS1, and DS2 enhancers as expected. This was also unchanged (not statistically increased) with PTH treatment. Like VDR recruitment in Fig. 5, A and B, pCREB recruitment was reduced overall, however there continued to be no change in the occupancy of pCREB after PTH treatment (C24-V1V2, Fig. 5C, lower track). Additionally, the peak of pCREB binding at the PRO region was greatly reduced. While there appears to be some dependence on the PRO VDREs for the binding of CREB in the DS1 and DS2 regions, this appears to be unrelated to the ability to suppress *Cyp24a1* expression.

### Synergistic response of FGF23 and 1,25(OH)_2_D_3_ on vitamin D metabolic genes

Thus far, our data point to a potential synergistic relationship between 1,25(OH)_2_D_3_ and FGF23’s actions in controlling vitamin D metabolism. Each hormone pushes *Cyp24a1* and *Cyp27b1* in the same direction (induce and suppress, respectively). To validate this synergy, we tested co-treatments of 1,25(OH)_2_D_3_ and FGF23 for gene expression differences. The C24-V1V2 and WT littermates were both treated with vehicle (Veh), 1,25(OH)_2_D_3_ (6 h), FGF23 (3 h), or the combination of 1,25(OH)_2_D_3_ and FGF23. To maintain, the time-based maximal responses (9,10,22), 1,25(OH)_2_D_3_ (10 ng/g or 0.1 ng/g bw) was dosed first into the mice for 3 h, followed by an injection of 50 ng/g bw FGF23 and mice were sacrificed 3 h later (6 h total time). The 10 ng/g bw dose of 1,25(OH)_2_D_3_ is a supraphysiologic dose of 1,25(OH)_2_D_3_ in the animal and may be the maximal response in the animal. Therefore, we also conducted this experiment at an additional dose of 1,25(OH)_2_D_3_ that was 100-fold less at 0.1 ng/g bw. Since FGF23 only induces *Cyp24a1* in the kidney, we only examined the kidney *Cyp24a1* and *Cyp27b1* expression after injection. As can be seen in Fig. 7A, the *Cyp24a1* expression is induced by FGF23 and 1,25(OH)_2_D_3_ at both the 0.1 and 10 ng/g bw doses. Additionally, the co-treatments of FGF23 with either 1,25(OH)_2_D_3_ concentration resulted in a significant increase in *Cyp24a1* expression versus individual treatments. The expression of *Cyp27b1* was robustly suppressed by the 10 ng/g bw dose of 1,25(OH)_2_D_3_ therefore, it was difficult to detect the further suppression of *Cyp27b1* as the values approached the limits of detection by qPCR (Fig. 7B). These data demonstrate a synergistic activation of *Cyp24a1* and suppression of *Cyp27b1* with 1,25(OH)_2_D_3_ and FGF23 together.

**Figure 7:**
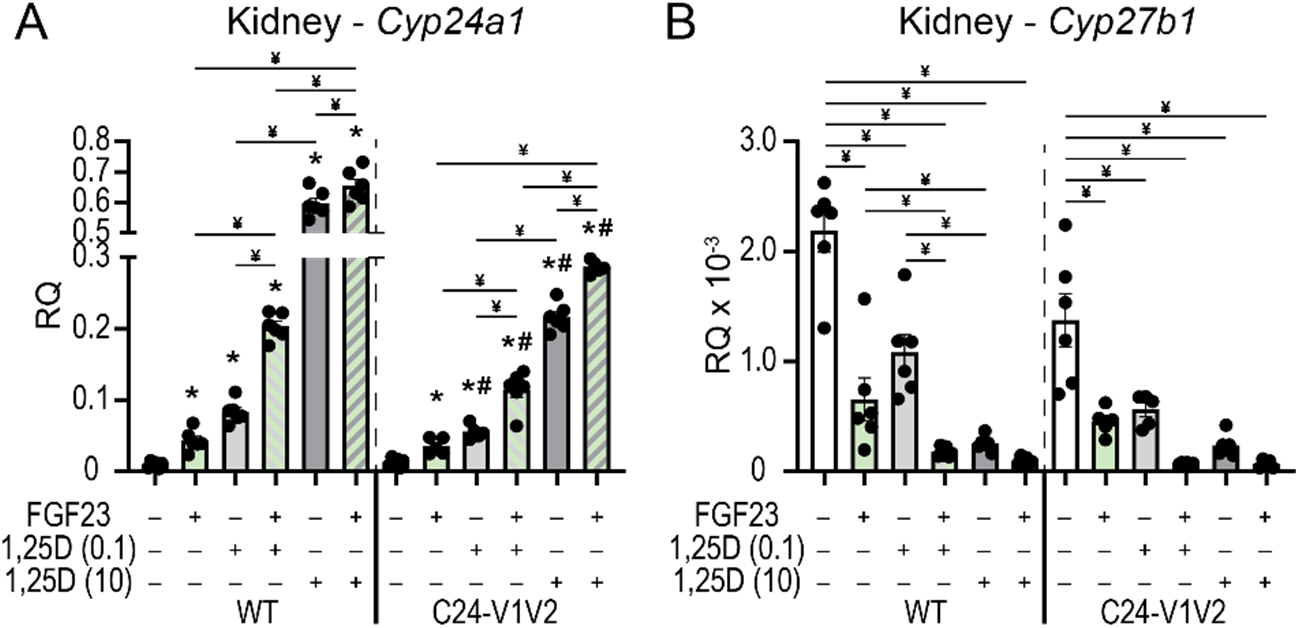
Synergistic actions of 1,25(OH)_2_D_3_ and FGF23 in the regulation of *Cyp24a1*. Gene expression was performed in WT and C24-V1V2 mice injected with ethanol vehicle (Veh), 50 ng/g bw FGF23 (3 h), 0.1 ng/g bw 1,25(OH)_2_D_3_ (6 h), 10 ng/g bw 1,25(OH)_2_D_3_ (6 h), FGF23 + 0.1 ng/g bw 1,25(OH)_2_D_3_, or FGF23 + 10 ng/g bw 1,25(OH)_2_D_3_ for kidney *Cyp24a1* (A) and *Cyp27b1* (B). Data are displayed as relative quantitation (RQ) against *Gapdh*. All groups, n ≥ 6. 2-way ANOVA with tukey post-tests comparing all treatments and genotypes was performed. ^*^, p < 0.05, treatment vs Veh; #, p < 0.05, C24-V1V2 vs WT; ¥, p < 0.05, selected pairs as indicated.

### PTH inhibits the ability of FGF23 to stimulate Cyp24a1 expression in the kidney

As identified above, FGF23 not only stimulates *Cyp24a1* expression in the kidney via the *Cyp24a1* DS1, but this hormone also synergizes with 1,25(OH)_2_D_3_ as seen in Fig 7, to increase *Cyp24a1* expression in that tissue via the DS2. Interestingly, in a further hormonal interaction, PTH also prevents FGF23 but not 1,25(OH)_2_D_3_ from inducing *Cyp24a1*, as seen in Fig 8 and in our previous work (9,22). Thus, under high PTH conditions, as seen in un-rescued *Cyp27b1*-KO and M1/M21-DIKO mice with high PTH levels (9,22), while *Cyp24a1* remains induced by treatment with 1,25(OH)_2_D_3_, FGF23 is no longer able to upregulate this gene in the kidney (Fig. 8). These results suggest that in addition to regulation, FGF23 and PTH alter the turnover rate of 1,25(OH)_2_D_3_ in this tissue. Given the location of the opposing actions of FGF23 and PTH on *Cyp24a1* expression in the same region (DS1) independent of 1,25(OH)_2_D_3_ induction (DS2) in the same region, DS1 may be an influential regulatory switch for *Cyp24a1* expression in the kidney.

**Figure 8:**
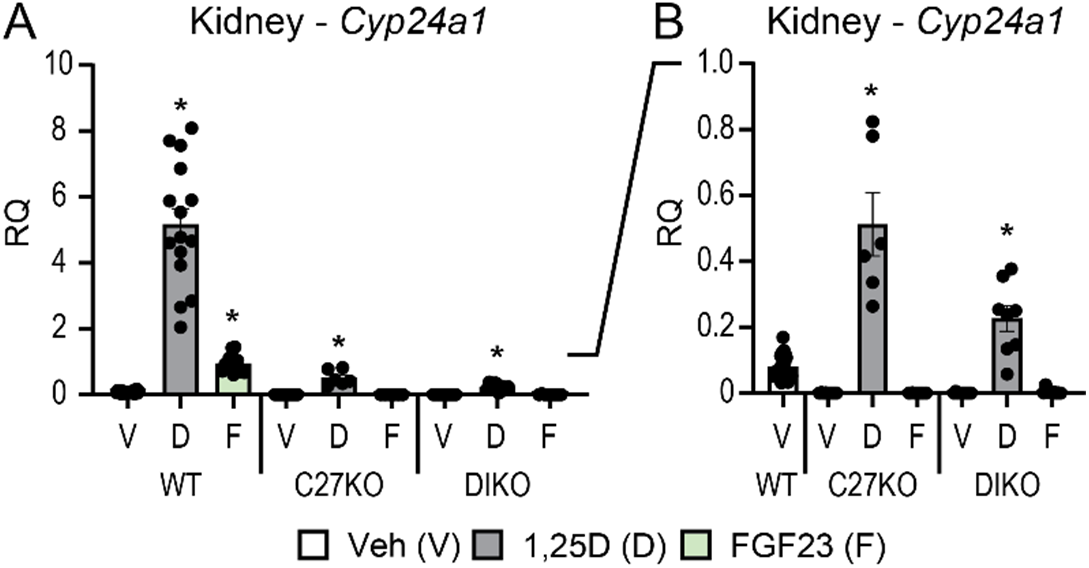
PTH inhibition of *Cyp24a1* activity in the kidney. Gene expression was performed in WT, *Cyp27b1*-KO (C27KO), and M1/M21-DIKO (DIKO) mice injected with 10 ng/g bw 1,25(OH)_2_D_3_ (6 h), 50 ng/g bw FGF23 (3 h), or PBS/ethanol vehicle (Veh) for kidney *Cyp24a1*. A, full scale data (0-10.0 RQ). B, subset of data in (A) with scale range from 0 – 1.0 RQ. Data are displayed as relative quantitation (RQ) against *Gapdh*. WT (Veh, 1,25(OH)_2_D_3_, FGF23) n ≥ 10; *Cyp27b1*-KO (Veh, 1,25(OH)_2_D_3_, FGF23), n ≥ 5; M1/M21-DIKO (Veh, 1,25(OH)_2_D_3_, FGF23), n ≥ 6. 2-way ANOVA with tukey post-tests comparing both treatment and genotype was performed. ^*^, p < 0.05, treatment vs Veh.

## Discussion

Expression and regulation of *Cyp24a1* in the mouse is essential for mineral homeostasis and vitamin D metabolism. Mice that lack global *Cyp24a1* expression (*Cyp24a1*-KO) have hypervitaminosis D, hypercalcemia, and secondary hyperparathyroidism that can lead to death (34,35), however a renal-specific deletion has not been created. Complex endocrine hormone regulation of *Cyp24a1* involves a combination of the promoter proximal enhancer and downstream enhancers for full activation (10,18). Our prior elimination of the DS1 region reduced basal expression of *Cyp24a1* and removed the majority of the regulation by FGF23 and PTH, thus assigning a tissue-specific role for the DS1 region (10). Furthermore, we previously found that the DS2 region bound VDR and was involved in 1,25(OH)_2_D_3_-mediated activation (10). Reporter assays, bacterial artificial chromosome (BAC) reporters, and common sense indicate that these near perfect consensus VDREs in the PRO region were important for *Cyp24a1* expression, however it was unknown how essential they were for activity considering the presence and activity of the DS regions (10,18). It also remained a mystery as to how these DS regions were able to affect transcription of *Cyp24a1* and if the PRO region was the connection point in three-dimensional structure to regulate *Cyp24a1*.

The VDRE sequences had been tested *in vitro* and had been found to activate reporter constructs in transfections (16,36), however these mutations were never made *in vivo*. It was unclear if one or both VDREs were needed *in vivo* for activation of *Cyp24a1*. Multiple VDREs had been discovered close to a number of genes and were found to assist in their regulation (19,37). One such example was intestinal regulation of the mouse calcium channel *Trpv6* in which there were not only multiple enhancers, but also multiple VDREs present that contributed to activation (38). With expanding genomic data sets for nuclear receptors, including our own work for VDR, these types of enhancers and multiple response elements were found to be more the rule than the exception. In fact, deletion of regulatory elements is often confounded by the number of putative elements that are discovered by computational methods, so much so, that it is easier to delete the enhancer in bulk to determine *in vivo* activity versus individual elements. For *Cyp24a1*, these VDREs were well-characterized *in vitro* so we used homology-directed recombination to delete the elements seamlessly and to ensure the genomic distance or sequence was not otherwise altered. We found that VDRE1 was the far more active VDRE as loss of VDRE2 alone (C24-V2) showed a rather mild change in gene expression compared to the C24-V1V2 mouse at both the kidney and intestine (Fig. 2). Despite their contributions to 1,25(OH)_2_D_3_-mediated expression, neither the C24-V1V2, nor the C24-V2 mouse had any loss of basal activity. This eliminates VDR binding in the PRO region as a basal driving enhancer unlike the DS1 where the basal levels of *Cyp24a1* were reduced with DS1 deletion (10). The DS1 is kidney-specific unlike the ubiquitous nature of both the PRO and DS2 regions, and the kidney has the highest expression levels of *Cyp24a1*, therefore it is likely that DS1 alone drives the tissue-specific basal expression of *Cyp24a1*.

Despite the lack of baseline drop of *Cyp24a1* expression in either the C24-V1V2 or C24-V2 mice, there was a drop in the baseline of *Cyp27b1* in the kidney. We saw a compensatory baseline drop in *Cyp24a1* when we mutated the enhancers of the *Cyp27b1* gene (9,22), and additionally, we see the *Cyp27b1* baseline drop when we delete the DS1 region of *Cyp24a1* (10). These activities are compensatory in response to the shifts in Ca and P in the blood and in turn the PTH and FGF23 in circulation. The homeostatic system attempts to correct these levels and put the mouse back at equilibrium. In some animal models, the deficiencies are too extreme and mineral loss too great, so these animals are incapable of achieving homeostasis without dietary interventions such as the global *Cyp27b1*-KO mouse and the M1/M21-DIKO model of disrupted renal vitamin D metabolism (9). However, in our C24-V1V2 and C24-V2 mice, it appears from their WT serum phenotype (normal Ca, P, PTH, iFGF23, Fig. 1) that these animals achieved mineral homeostasis. Interestingly though, even with normal serum parameters, these animals both have a mild elevation of 25(OH)D_3_ in the blood. This indicates that either or both CYP27B1 and CYP24A1 enzymes have less activity, as we observe a substrate (25(OH)D_3_) excess with reduction of enzymatic activity. While the 25(OH)D_3_ levels are increased, so too are the 24,25(OH)_2_D_3_ levels (∼16 ng/mL to 20 ng/mL 25(OH)D_3_ and ∼ 6 ng/mL to 10 ng/mL 24,25(OH)_2_D_3_, Fig. 1C). These data indicate that CYP24A1 activity may very well be normal and perhaps the CYP27B1 lowered expression was causing the increase of 25(OH)D_3_ substrate. This was supported by the elevated levels of 24,25(OH)_2_D_3_ and the production of the 3-epi-25(OH)D_3_ product. The compensatory reduction of *Cyp27b1* expression appeared abnormal given the WT serum profiles. Despite the lower *Cyp27b1* expression, the C24-V1V2 animal was making enough 1,25(OH)_2_D_3_ to remain normal and healthy, so the homeostatic system was functioning as intended.

After the PRO VDRE mutations of the C24-V1V2 mouse, we found that the levels of VDR occupancy were dramatically decreased but not completely eliminated at the PRO region. An *in silico* analysis of the mutated sequences did not produce many high-scoring putative VDREs (data not shown). This could simply be a low-ranking VDRE still binding a small amount of VDR, or more interestingly, could be residual binding from the DS1/2 regions being captured by the ChIP crosslinking in close proximity. The retained VDR binding at the PRO was also not responsive to 1,25(OH)_2_D_3_ treatment (Fig. 5, A and B, no increase of blue). Distal enhancers are often brought into close proximity via three-dimensional structures (39), so perhaps we have severed most of the connection point with elimination of the VDREs, but not all. These data are supported by the fact that binding of VDR was also reduced in the DS1 and DS2 regions (Fig. 5). VDR occupancy was still increased in the DS2 region, but to much lower amplitude. This also raised the question of DS1 connections to the PRO to aid in transcription and if VDR at the PRO VDREs was involved in that relationship. We found that like VDR, the existing CREB binding to the *Cyp24a1* PRO was also eliminated with the mutation of the VDREs as were the downstream DS1/2 CREB occupancies. However, this had no impact on *Cyp24a1* suppression by PTH (Fig. 4A). Therefore, CREB could be involved in the suppression mechanism through recruitment of chromatin suppressors and histone deacetylases, however, it may be that an additional unknown factor is facilitating this suppression by removal from the genome. This would also support the removal of coactivators we observed in our recent study of the rapid actions of PTH on *Cyp27b1* and *Cyp24a1* actions (21). We found CBP, CRTC2, SRC1, and others removed from the genome after PTH treatment, which was not the case for CREB (21). We hypothesize that this unknown factor might also counteract the FGF23-mediated increases in *Cyp24a1* given the co-localization of activities to the DS1 region where both PTH and FGF23 actions reside. The actions of FGF23 are affected by the mutation of the PRO VDREs. There was a reduction of the induction of *Cyp24a1* with FGF23 treatment by approximately 50% (Fig. 4A). This may indicate that the FGF23 actions are tied together with VDR/1,25(OH)_2_D_3_ in at least the promoter region, more so than the PTH actions. We found a synergistic response of FGF23 and 1,25(OH)_2_D_3_ co-treatments on *Cyp24a1* that would support such a relationship (Fig. 7). While the lack of transcription factor for FGF23 actions hinders an extended analysis of these activities, it is certainly possible that there is an interaction between this unknown TF and VDR in the control of *Cyp24a1*. Together, this putative factor could both be involved in FGF23 activation through genomic recruitment and suppression via PTH by genomic removal to fit the model of activities we observe.

Given these activities, we believe the DS1 region acts as a molecular switch for *Cyp24a1* activity and thus for vitamin D metabolism. FGF23 flips this switch to “on” and allows expression of *Cyp24a1* to proceed, PTH acts as the brake and flips the switch to “off”. This switch operates in the kidney, under normal conditions, with micro-adjustments to keep the homeostasis in balance. The switch is then flipped under stressors to serum Ca (PTH) or P (FGF23) levels. This regulation influences the turnover of 1,25(OH)_2_D_3_ by virtue of the reciprocal control of *Cyp27b1* expression and the reduced degradation of 1,25(OH)_2_D_3_ via the CYP24A1 enzyme. This has particular importance in disease states like chronic kidney disease, where first FGF23 increases which wastes phosphate, exacerbates disease, and drives down vitamin D metabolism by activation of *Cyp24a1* and suppression of *Cyp27b1* (40,41). This is then followed by an elevation in PTH causing a further dysregulation of vitamin D metabolism. When PTH is high, in a model like *Cyp27b1*-KO, *Vdr*-KO, or M1/M21-DIKO mice (Fig. 8), the switch is locked off and requires dietary intervention to reduce the PTH levels by either recovering serum Ca levels or increasing 1,25(OH)_2_D_3_ through exogenous means (8,9,22,30). Additionally, when PTH is high, the skeleton suffers tremendously, and this can be predicted by examining the ratio of 25(OH)D_3_ to 24,25(OH)_2_D_3_ along with the activity of *Cyp24a1* as we have previously modeled (30). In the C24-V1V2 and C24-V2 animals, PTH is not elevated, vitamin D metabolism is in balance, and the mice and their skeletons are normal.

In conclusion, we found that the VDREs at the promoter of *Cyp24a1* are necessary for full activation of *Cyp24a1*, however, the VDREs are not essential for mineral homeostasis and vitamin D metabolism. This work highlights the interconnected relationships between 1,25(OH)_2_D_3_, PTH, and FGF23 and the utilization of multiple interconnected enhancers. Furthermore, these data point to a potential FGF23-responsive factor that contributes to the baseline of *Cyp24a1* that may be connected to each of these hormones for ultimate control of vitamin D metabolism.

## Experimental Procedures

### Reagents

The following reagents were used for *in vivo* injections: 1α,25(OH)_2_D_3_ was obtained from SAFC Global (Madison, WI), Parathyroid Hormone (PTH, 1-84 human) was obtained from Bachem (H-1370.0100 Torrence, CA), mouse Fibroblast Growth Factor 23 from R&D systems (FGF23, 2629-FG-025, Minneapolis, MN). ChIP antibody for VDR (C-20, sc-1008, lot # H1216) was purchased from Santa Cruz Biotechnology, Inc. (Santa Cruz, CA). Phosphorylated-CREB (Ser133) (pCREB, 06-519, lot # 3460466) was purchased from EMD Millipore (Burlington, MA). Traditional genotyping PCR was completed with GoTaq (Promega, Madison, WI) and all real-time qPCR was completed with the StepOnePlus (Applied Biosystems, Foster City, CA) using TaqMan for gene expression assays (Applied Biosystems). Primers for genotyping and CRISPR were obtained from IDT (Coralville, IA).

### Animal Studies

C57BL/6 mice aged 8-9 w (The Jackson Laboratory, Bar Harbor, ME) were housed in high density ventilated caging in the Nutritional Sciences Animal Facility at the University of Wisconsin-Madison under 12-hour light/dark cycles at 72°F and 45% humidity. All mice used in this study were maintained on a standard rodent chow diet (2020 diet, Envigo). All animal studies were reviewed and approved by the Research Animal Care and Use Committee of University of Wisconsin-Madison under protocol A005478. Animals were subjected to intraperitoneal injection of 10 ng/g body weight (bw) 1,25(OH)_2_D_3_ (in propylene glycol), 230 ng/g bw PTH (1–84) (in phosphate buffered saline (PBS)), 50 ng/g bw FGF23 (in PBS), or vehicle (EtOH or PBS). Animals were sacrificed and tissues collected at times indicated in each legend for ChIP and gene expression. Unless otherwise indicated, all experiments were conducted with equal numbers of males and females (n ≥ 6). Data were reported as mixed, as no differences were found between sexes.

### CRISPR mouse generation

The guides used for CRISPR-Cas9 editing were optimized for the least number of potential off-target sites and fewest sites in coding exons using the CRISPRko and CRISPick sgRNA Designer Tool from the BROAD Institute based on (42). Guides were prepared as previously reported (9,22). Guides and primers with genotyping schema are listed in Table 1. mC24-Pro-G5: GGGGACCGACTGGCAAGGAT – GGG and mC24-Pro-G6: CGCTTGCACAATCGCCACTC – AGG were used to target the promoter region. The ssODN megamer was purchased from IDT (Coralville, IA). The mixture of 50 ng/µL of the produced RNA guides, 50 ng/µL ssODN, and 40 ng/µL of Cas9 protein in injection buffer (5mM Trizma Base, 5mM Tris HCL, 0.1mM EDTA, pH 7.4) was injected into the pronucleus of one day fertilized embryos isolated from hyper-ovulating female C57BL/6 mice as described previously (43) and implanted into recipient females by the University of Wisconsin Madison Biotechnology Genome Editing and Animal Models Core. All CRISPR generated mice were backcrossed into C57BL/6J mice for 4 generations before experiments were performed to reduce any potential off-target mutations. DNA sequences were also analyzed and compiled from ChIP input DNA to ensure fidelity of desired genomic mutations.

**Table 1:**
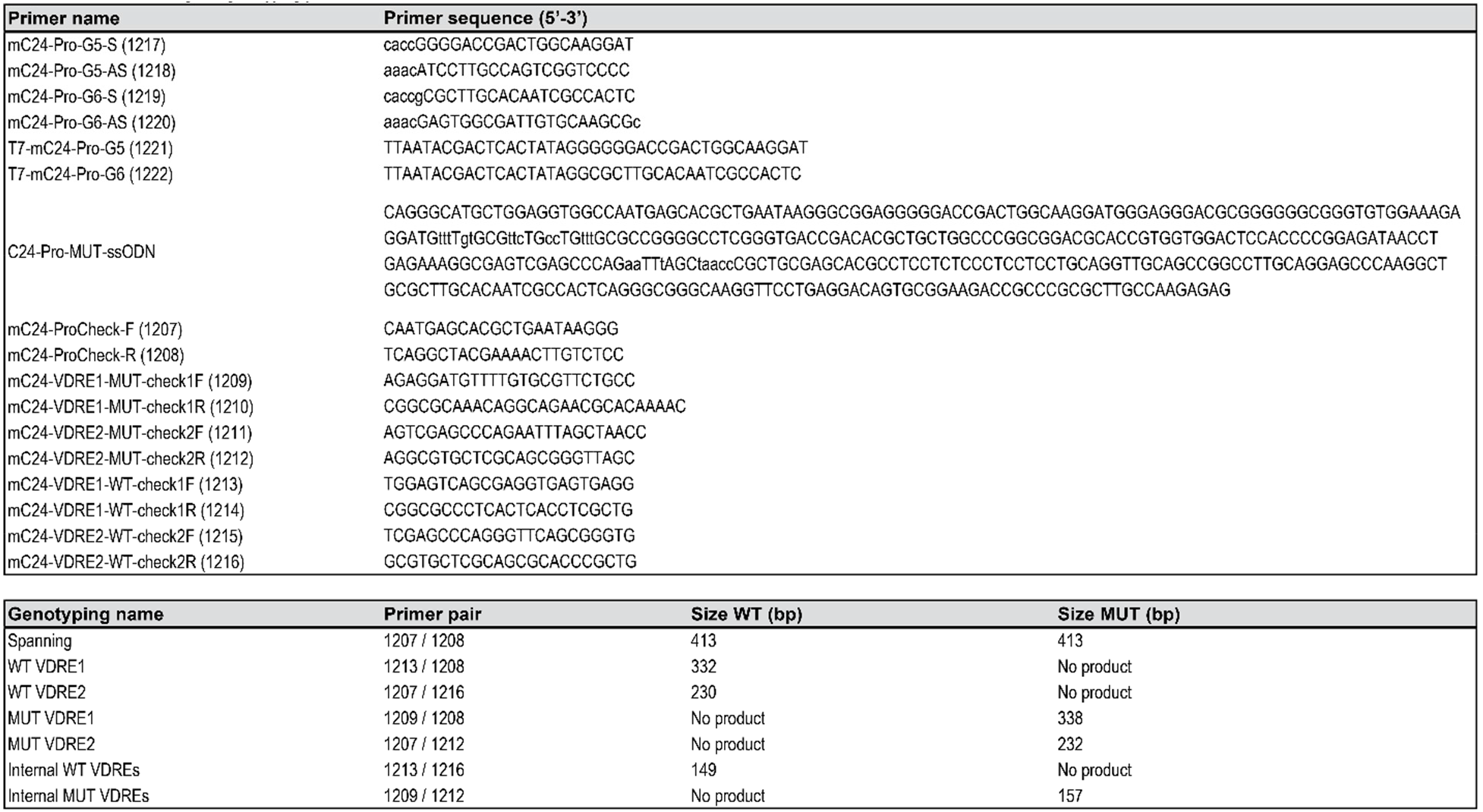
CRISPR cloning and genotyping primers.

### Blood Chemistry

Cardiac blood was collected at the time of sacrifice. Collected blood was split into serum or EDTA-treated plasma, incubated at room temperature for 30 min followed by centrifugation at 6,000 rpm for 12 min (x2) to obtain serum or EDTA-plasma(22). Serum calcium and phosphate levels were measured using QuantiChrom™ Calcium Assay Kit (#DICA-500, BioAssay Systems, Hayward, CA) and QuantiChrom™ Phosphate Assay Kit (#DIPI-500, BioAssay Systems, Hayward, CA)(22). Circulating intact FGF23 and PTH were measured in EDTA plasma via a Mouse/Rat FGF-23 (Intact) ELISA Kit (#60-6800, Immutopics, San Clemente, CA) and a Mouse PTH (1-84) ELISA kit (#60-2305, Immutopics), respectively (22).

### Gene Expression

Dissected tissues were frozen immediately in liquid nitrogen and stored at -80°C. Frozen tissues were homogenized in Trizol Reagent (Life Technologies) and RNA was isolated as per the manufacturer’s instructions. 1 µg of isolated total RNA was DNase treated, reverse transcribed using the High-Capacity cDNA Kit (Applied Biosystems), and then diluted to 100 µL with RNase/DNase free water. qPCR was performed using primers specific to a select set of differentially expressed genes by TaqMan analyses. TaqMan Gene Expression probes (Applied Biosystems) used were *Cyp27b1* (Mm01165918_g1, FAM), *Cyp24a1* (Mm00487244_m1, FAM), *Vdr* (Mm00437297_m1, FAM), *Trpv6* (Mm00499069_m1, FAM), and normalized to *Gapdh* (Mm99999915_g1, VIC).

### ChIP followed by Sequencing (ChIP-seq)

Chromatin Immunoprecipitation (ChIP) was performed using antibodies listed in Reagents. ChIP was performed as described previously in triplicate (both male and female mice were used, n=3 per treatment) with several modifications (9,10,21). Due to library loss, the pCREB vehicle treated mice were n=2. The isolated DNA (or Input DNA acquired prior to precipitation) was then validated by quantitative real time PCR (qPCR) and further prepared for ChIP-seq analysis. ChIP-seq libraries were prepared as previously described (44,45) with the following exceptions: ChIP-seq libraries were prepared using the NEBNext Ultra II DNA kit (NEB, #E7645S) with the NEBNext Multiplex Oligos for Illumina (NEB, #E6440S) according to the manufacturer’s protocols with library concentrations measured by Qubit (Life Technologies) and library size distribution by Bioanalyzer 2100 (Agilent). Libraries were sequenced on a NovaSeqX (Azenta/Genewiz, Waltham, MA). Paired end, 150 bp sequencing with a target of 30+ million reads was performed. ChIP-sequencing files were processed by HOMER, EdgeR, and DESeq2 and the raw and processed data (bigWig files) have been deposited in the GEO archives (mm9 genome, see Data availability). The bigWig files for replicates were averaged using wiggletools mean function. All ChIP data were normalized to 10 million reads and to Input DNA for display in the UCSC Genome Browser as overlaid track hubs.

### Statistical Evaluation

Data were analyzed using GraphPad Prism 10.3 software (GraphPad Software, Inc., La Jolla, CA) and in consultation with the University of Wisconsin Statistics Department. All values are reported as the mean ± standard error (SEM) and differences between group means were evaluated using One-Way ANOVA, Two-Way ANOVA, or Student’s *t*-test as indicated in the figure legends.

## Data availability

Raw and processed ChIP-seq data have been deposited in the GEO archives at GSE275331. The kidney WT Input are deposited in GSM3898546 and the pCREB WT samples in GSE206777. All remaining data have been included in the manuscript, the supplements, or are freely available upon request.

## Acknowledgements

We thank members of the Meyer Laboratory for their contributions during manuscript preparation. We also thank Drs. Glenville Jones and Martin Kaufmann for the measurements of the vitamin D metabolites by LC-MS/MS.

## Author contributions

Conceptualization, M.B.M.; Investigation, M.B.M., S.M.L., J.M.T., S.R.C., M.K.; Writing – Original Draft, M.B.M.; Writing – Review & Editing, M.B.M., J.W.P., S.M.L., J.M.T., S.R.C., G.J., M.K.; Data Curation, M.B.M.

## FOOTNOTES

## Abbreviations used

1,25(OH)_2_D_3_: 1,25 dihydroxy vitamin D
FGF23: fibroblast growth factor 23
PTH: parathyroid hormone
Ca: calcium
P: phosphate
pCREB: phosphorylated cAMP response element-binding protein
CBP: CREB-binding protein
PKA: protein kinase A
C24-PRO: *Cyp24a1* promoter proximal region
C24-V1V2: *Cyp24a1* promoter mutations in VDRE1/AP-1 and VDRE2
C24-V2: *Cyp24a1* promoter mutations in VDRE2 only
DS1: DownStream 1 region of *Cyp24a1*
DS2: DownStream 2 region of *Cyp24a1*
M1: *Mettl1* intronic enhancer of *Cyp27b1*
M21: *Mettl21b* intronic enhancer of *Cyp27b1*.

